# First high-fidelity scaled 3D-printed models of insect tympanic membrane and acoustic trachea preserving their acoustic function

**DOI:** 10.1101/2025.10.24.684398

**Authors:** Md Niamul Islam, Fabio A. Sarria-S, Fernando Montealegre-Z

## Abstract

Miniature dual-input hearing in katydids underpins communication and bat evasion, yet its microscale anatomy hinders acoustic studies. With µCT imaging, AI-assisted segmentation and multi-material 3D printed assembly, scaled copies of high-fidelity pinna-tympanum assembly and a complete acoustic trachea of the neotropical katydid *Copiphora gorgonensis* were fabricated. Flexible TPU membranes reproduce similar tympanal vibrations compared to actual insect and pairing with rigid PLA pinnae mimicked the outer-ear motion, providing ultrasonic gain at 70–110 kHz matching *in vivo* bat-detection bands. Separately, the pressure mapping of the scaled acoustic trachea confirms the spiracle as a spectral filter and the exponential canal as a 17–21 dB amplifier, in line with simulations, preserving the 1.3 cycle phase shift seen at 23 kHz in living insects. These matching results justify the use of scaled biomimicking replicas as reusable, 3Rs-aligned substitutes for living insect acoustic studies in search bioinspired applications.

## 1. Introduction

Acoustic localisation is a fundamental survival trait: predators must pinpoint prey, prey must track predators, and both must identify conspecifics [1]. Most vertebrates solve this task with binaural cues — inter-aural time differences (ITDs) and inter-aural level differences (ILDs) [2]. For miniature animals, however, head width quickly approaches a fraction of the relevant wavelength, compressing ITDs to microseconds and ILDs to fractions of a decibel [3]. Katydids (or bush-crickets: Orthoptera, Tettigoniidae), a clade exceeding 8,300 described species [4], overcome this scaling problem by relocating their ears to the forelegs. Separating the pair of ears by the width of the body, rather than the head, enlarges path-length disparity and restores usable phase lags even for ultrasonic signals emitted in courtship or predation [5,6].

The katydid also remains unique among invertebrates for developing a hearing mechanism similar to that of mammals [7,8]. Each ear contains two thin tympana situated on the proximal tibia and supplied by dual acoustic routes. External sound acts directly on the membrane surface, whereas internal sound enters a thoracic spiracle, propagates through an exponential acoustic trachea and impinges upon the inner membrane face [9,10]. In *Copiphora gorgonensis*, the exponential horn-shaped trachea boosts pressure by ∼15 dB and slows phase by approximately 25 % relative to the external path, generating a pressure-difference receiver [9]. More than 65 % of katydids add a third refinement: cuticular pinnae fold over the tympana, creating Helmholtz-like cavities that deliver an additional 20–30 dB gains above 60 kHz, matching the echolocation band of gleaning bats [11,12]. A fluid-filled cavity covering the *crista acustica* act as a dispersive travelling-wave analyser, mirroring the tonotopic processing of the vertebrate cochlea [13].

Direct experimental work at the micro-scale of the katydid ear presents a significant challenge. The tympanic membranes are delicate, while the acoustic trachea is buried deep within the tibia [14]. Thus, non-invasive techniques are employed, laser-doppler vibrometry of intact specimens [15] and finite-element models built from µCT volumes [16–18], which also limits the functional insights of ear mechanics. However, a better alternative is experimenting on scaled replicas fabricated via 3D printing (additive manufacturing). This valuable manufacturing process is capable of accurately replicating the complex anatomy of insect auditory structures. Scaled replicas are effective for acoustic experiments because sound-structure interactions depend on the ratio between the structure’s size and the wavelength of sound, rather than their absolute values. For example, if the ear model is enlarged ten times (10×) and tested at sound frequencies ten times lower (1/10×), these dimensionless relationships remain unchanged. As a result, wave propagation, resonance, and vibration patterns occur in the same relative way as in the natural ear. Using this phenomena, existing studies used 3D printing to replicate the function of the pinnae [11] and produce simplified insect ear models with flexible membrane material [19].

However, the precise fabrication of high-fidelity fully functional katydid outer ear, flexible tympanic membranes with the rigid cuticle and pinnae remains a significant challenge, as demonstrated in recent multimodal additive manufacturing of tympanic membranes replicating their acousto-mechanical performance [20]. The complex structural geometry and differences in the stiffness of printing materials, adhesion limitations, and segmentation accuracy pose additional obstacles to creating biologically accurate scaled models [21]. Despite these challenges, achieving a fully functional 3D-printed outer ear and acoustic trachea offers considerable advantages, as it enables scalability for biophysical measurements and controlled manipulation of structural parameters such as horn flare, cavity volume, and membrane stiffness. Working with scaled-up replicas not only simplifies experimental procedures and visualization but also allows for future systematic testing of design variations to uncover the principles of frequency filtering and mechanical coupling [16]. These models provide a powerful platform for validating numerical simulations, optimizing structural performance, and guiding the development of bio-mimicking auditory systems [22].

In this study, we present the first scaled, 3D-printed multi-material functional replica of the outer ear model of the neotropical katydid *Copiphora gorgonensis*, integrating flexible tympana with rigid pinnae and the complete spiracle-to-tympanum acoustic trachea. Built from high-resolution µCT data, AI-assisted segmentation, and advanced 3D printing techniques. The promising bioacoustics response of the scaled models establishes a platform for systematic investigation of miniature filters, amplifiers and pressure-difference receivers, providing a tangible pathway from insect biomechanics to next-generation acoustic sensors. As additive manufacturing technologies continue to advance, with increasing printing resolution and multi-material precision, the same design principles can be scaled down to create miniature bio-inspired acoustic sensors, including MEMS-based devices [19,23,24]. Thus, 3D printing not only facilitates current research through accessible, controllable scaled replicas but also paves the way toward practical implementations of insect-inspired auditory mechanisms in compact sensing systems. Additionally, the approach supports the 3Rs principles (Replacement, Reduction, and Refinement) by reducing dependence on live insects, minimizing experimental variability, and improving reproducibility and control [25].

## 2. Methodology

### 2.1. Specimen

The specimens of *Copiphora gorgonensis* (Tettigoniidae: Copiphorini), which are endemic to Gorgona National Natural Park, Colombia, were imported to the United Kingdom in 2015 under a research permit issued by the Colombian Authority (DTS0-G-090 14/08/2014). These specimens were bred into ninth-generation captive colonies and maintained under carefully controlled laboratory conditions, including a temperature of 25 °C, relative humidity of 70%, and a 12:12 light-to-dark cycle. The colonies were provided with an *ad libitum* diet consisting of bee pollen, fresh apple, dog food pellets, and water to ensure optimal development and health. Following the completion of bioacoustics experiments, the live specimens were preserved in increasing concentration of ethanol-filled centrifuge tubes up to 100% (to reduce excess water loss and shrinkage) and subsequently stored at −22 °C in a dedicated freezer at the University of Lincoln for further analysis and imaging.

### 2.2. Micro-computed Tomography (µCT) scanning

Two preserved adults *Copiphora gorgonensis* specimens were imaged on a SkyScan 1172 µCT scanner (Bruker Corporation, Billerica, MA, USA), each tuned to a different structure. For the acoustic trachea, the anterior half of the first insect (head, prothorax and attached forelegs) was scanned without staining at 55 kV, 200 µA, 350 ms exposure and 0.1° rotation steps, giving a 5 µm voxel size that resolved the spiracle and acoustic trachea. The second specimen’s left foreleg was excised, stained for 24 h in 1 % iodine-ethanol to enhance membrane contrast, rinsed with ethanol and desiccated overnight in hexamethyldisilazane (HMDS) to minimise tympanal deformation. The dried leg was scanned at 50 kV, 180 µA, 300 ms exposure and 0.1° rotation steps, yielding 1.5 µm voxels that resolved the tympana together with their cuticular pinnae. The raw projection data were reconstructed into were exported as 16-bit TIFF orthogonal slices using NRecon software (v.1.6.9.18, Bruker Corporation, Billerica, MA, USA).

### 2.3. Deep (AI) semantic segmentation and volumetric reconstruction

Semantic segmentation of the insect outer ear was performed using the deep learning module in Dragonfly 3D software (Object Research Systems, Montreal, Canada), utilising a U-Net convolutional neural network architecture for iterative model training and refinement (Fig. 1) [26]. Semantic segmentation was employed instead of purely manual segmentation due to its significant advantages in accuracy, efficiency, and reproducibility. Manual segmentation of complex biological structures such as the tympanic membranes and acoustic trachea is extremely time-consuming and prone to user bias, as the boundaries between cuticle, air spaces, and soft tissue are often difficult to distinguish in µCT data. In contrast, deep learning-based segmentation using the U-Net architecture allows the model to learn subtle textural and intensity differences across large image datasets, enabling consistent and precise identification of anatomical features. This automated approach ensures high-fidelity reconstruction of the intricate 3D geometry of the ear, capturing realistic anatomical variations rather than relying on idealized or oversimplified models, which is essential for producing accurate bio-physical and acoustic simulations.

**Fig. 1.**
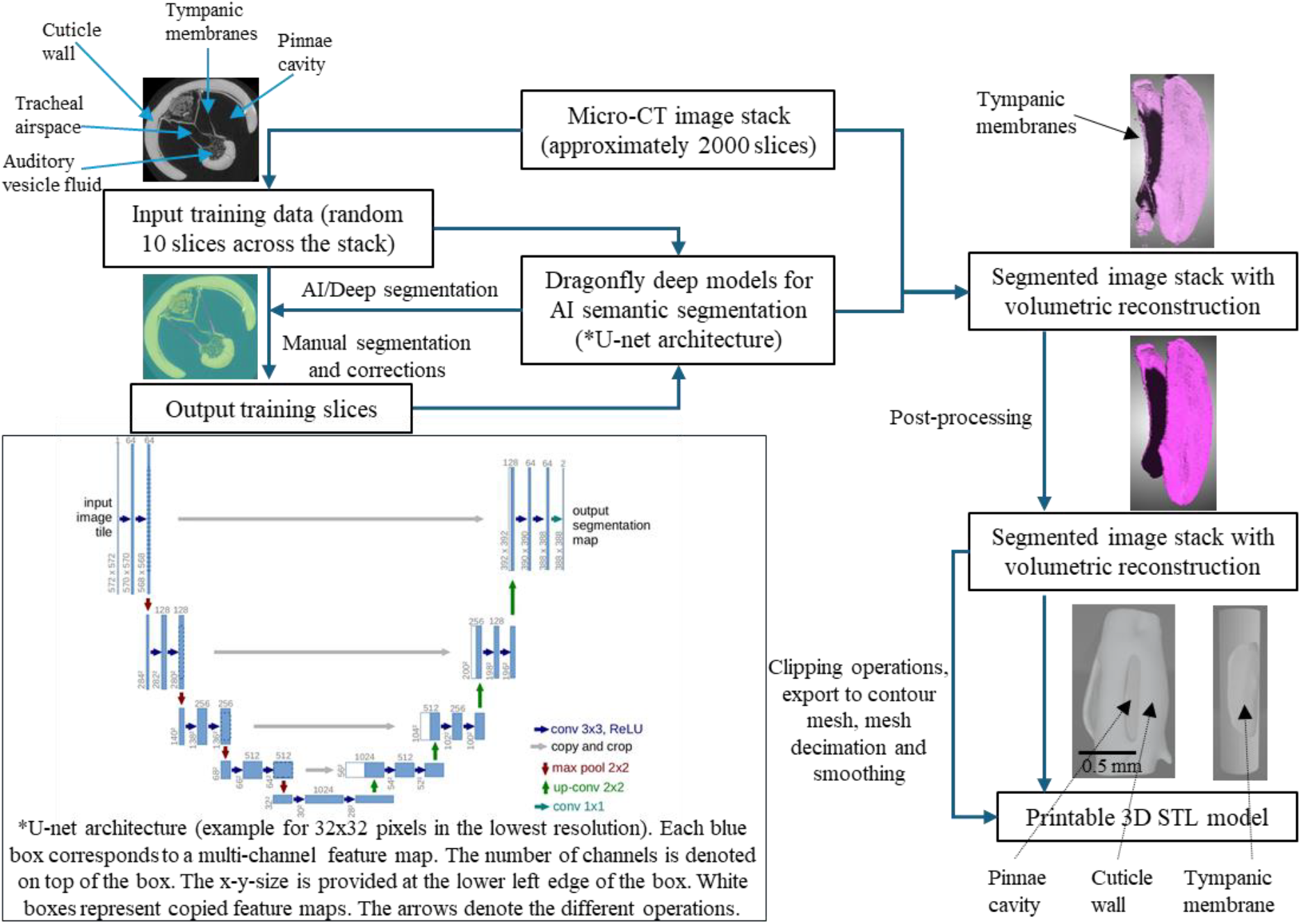
Deep learning-based semantic segmentation pipeline of the katydid outer ear for 3D reconstruction and fabrication. A U-Net convolutional neural network was trained on annotated µCT slices to segment the outer ear structures, including the cuticle, tympana, and air spaces. The model was refined over multiple iterations and applied to the full dataset to generate volumetric segmentations. These were processed to isolate the tympana and cuticle, enabling export as high-fidelity STL files for multi-material 3D printing.

For tympanic membrane segmentation, approximately 2,000 high-resolution µCT slices of the insect’s outer ear were imported into the software, from which ten slices were randomly selected to create a representative training subset, referred to as *Input Slice V1*. These slices were manually segmented to define multiple regions of interest (ROIs), including the cuticle, tympana, and air spaces, producing the corresponding labelled dataset, *Output Slice V1*. This dataset served as the ground truth for initial model training.

The U-Net model was first trained using *Input Slice V1* and *Output Slice V1*. To evaluate and improve model accuracy, a new set of ten slices (*Input Slice V2*) was randomly extracted from the full dataset and segmented using the trained model. The output segmentations (*Output Slice V2*) were manually reviewed and corrected to rectify classification errors. These corrected slices were then used to retrain the model. This process of prediction, correction, and retraining was repeated through three further iterations (up to *Input/Output Slice V5*), progressively enhancing model performance. After four training cycles, the model achieved accurate classification of anatomical structures with minimal need for manual correction and was applied to segment the entire volumetric dataset automatically.

After the segmentation of the outer ear, the tympana were isolated from the surrounding cuticle to enable material-specific 3D printing. A virtual cylindrical tool was precisely aligned with the tympanal region in the segmented volume to define the separation boundary. A clipping operation was performed to partition the flexible tympanal membranes from the rigid cuticle structures. This operation generated two physically distinct models for printing: a flexible tympanum and a rigid supporting cuticle. These were later assembled into a single anatomical model, faithfully replicating the geometry and functional differentiation of the insect’s outer ear.

A similar deep learning approach was employed to segment the acoustic tracheal system, using approximately 1,500 µCT slices. The U-Net model was trained to distinguish the air-filled tracheal lumen from surrounding tissues, enabling accurate reconstruction of the acoustic trachea.

### 2.4. 3D printing of segmented structures

Studies on the mechanical properties of insect cuticles have shown that their stiffness varies widely, with soft cuticle ranging from approximately 1 kPa to 50 MPa, while sclerotized (hardened) cuticle can reach values between 1 GPa and 20 GPa [27]. Based on this, two printing materials were selected: polylactic acid (PLA) and thermoplastic polyurethane (TPU), manufactured by Bambu Lab and procured from AdditiveX. PLA, with a Young’s modulus of about 2.7 GPa, was used to represent the rigid cuticular structures of the outer ear, providing the necessary stiffness and structural stability. In contrast, TPU (TPU 95AH), with a Young’s modulus of around 9.8 MPa, was chosen to replicate the flexible tympanic membranes, allowing realistic vibration and deformation under sound excitation similar to the natural tympanum. Additionally, models with opposite material combinations (PLA tympana and TPU cuticle) were also printed to compare how differences in material stiffness influence the overall acoustic and mechanical behaviour of the structure. PLA was also selected for the fabrication of the scaled acoustic trachea, as atomic force microscopy (AFM) nanoindentation measurements of the native tracheal wall revealed a broad distribution of elastic moduli, with a mean of 5,200 MPa and a median of 1,030 MPa, indicating a predominantly stiff structural profile [28].

The outer ear and acoustic trachea were 3D printed using the fused deposition modelling (FDM) process on a Bambu P1S printer (Bambu Lab, Shenzhen, China) operated via Bambu Studio software. The katydid outer ear model was enlarged 15 times, while the acoustic trachea was enlarged 20 times to fabricate scaled physical replicas suitable for acoustic experimentation (Fig. 2). The default printer settings, based on the drop-down material profiles in Bambu Studio, were used for most parameters. A few key settings were adjusted to improve print quality and structural accuracy: a 0.2 mm nozzle diameter, 0.06 mm layer height, nozzle temperature of 230 °C, and bed temperature of 30 °C were applied for both PLA and TPU structures. For the outer ear models, a 100% infill with concentric layering was used to ensure full rigidity, whereas the acoustic trachea was printed without infill, using a 0.25 mm shell thickness to maintain the hollow internal cavity essential for acoustic functionality.

**Fig. 2.**
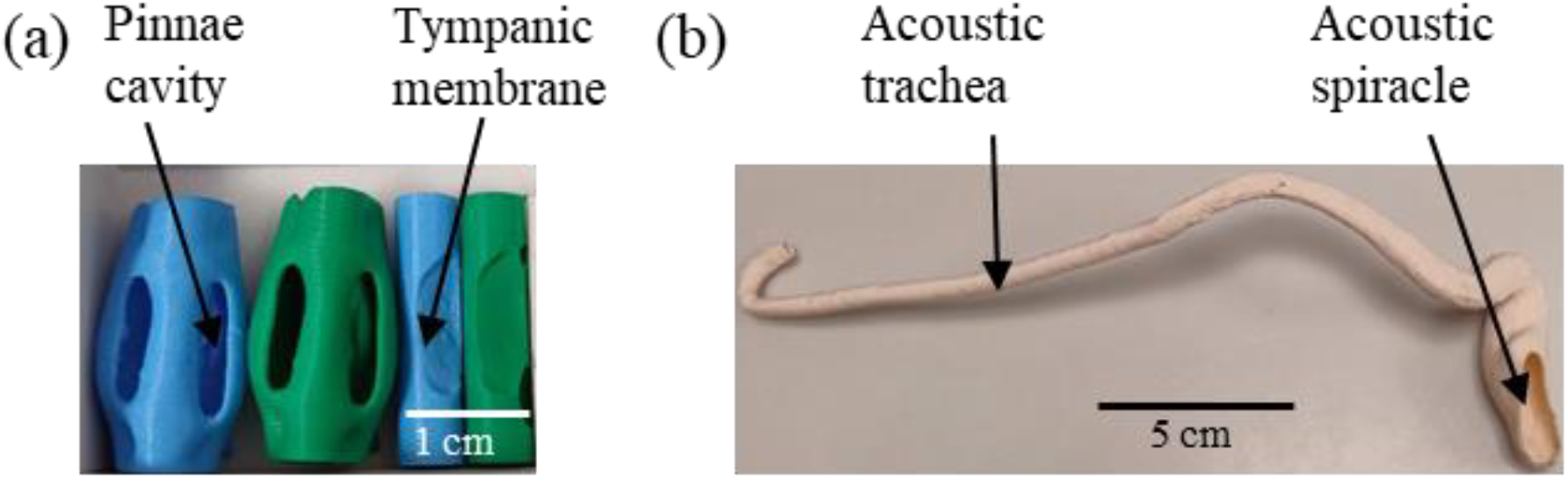
3D-printed models of the katydid outer ear and acoustic trachea. (a) 15× scaled outer ear components printed using FDM and SLA methods. PLA (green) and TPU (blue) structures were fabricated separately and assembled for acoustic testing. (b) 20× scaled acoustic trachea printed in PLA using FDM. Only the outer wall was printed to preserve the hollow lumen, replicating the native tubular geometry essential for acoustic functions.

### 2.5. Laser Doppler Vibrometry (LDV) on scaled tympana of the outer ear model

LDV experiments were conducted in an acoustic chamber designed to minimise external noise and vibrations (Fig. 3). All equipment and samples were placed on a vibration isolation air table to reduce measurement noise. The samples were mounted on a beam supported by a magnetic stand and fixed in position using adhesive putty. The orientation of the samples was adjusted to focus the laser beam from the scanning vibrometer (Polytec PSV-500 scanning head; Waldbronn, Germany) onto the tympanic membrane.

**Fig. 3.**
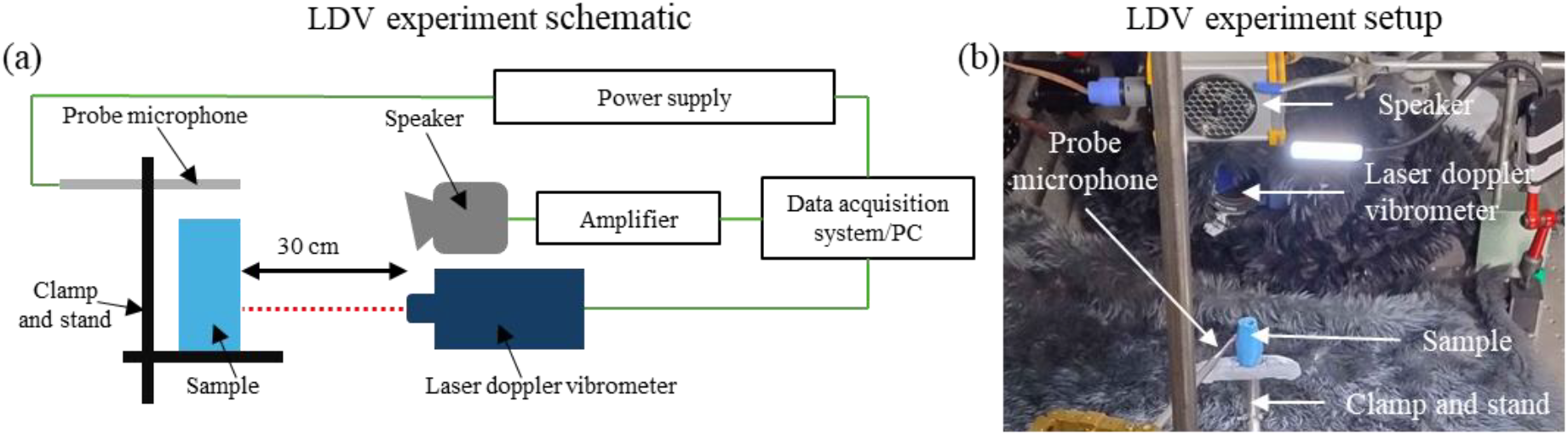
Laser Doppler vibrometry setup for vibration analysis of 3D-printed insect outer ear models. (a) Schematic of the experimental setup showing a speaker delivering sound stimuli to the sample, with vibration velocity measured by a laser Doppler vibrometer (LDV) and sound pressure recorded by a probe microphone. (b) Experimental configuration highlighting key components, including the speaker, LDV head, probe microphone, and 3D-printed sample mounted on a clamp and stand.

Avisoft ultrasonic speaker Vifa with SPEAKON connector (Avisoft-UltraSoundGate; Glienicke/Nordbahn, Germany), positioned 30 cm from the samples and secured on a magnetic stand, was used to deliver acoustic stimuli. The loudspeaker was connected to a portable ultrasonic power amplifier (Avisoft-UltraSoundGate; Glienicke/Nordbahn, Germany) to ensure controlled sound delivery. A 1/8″ precision pressure microphone (Bruel & Kjaer, 4138; Nærum, Denmark) was placed adjacent to the samples, facing the speaker and connected to a power supply (GRAS 12AA 2-Channel Power Module with gain and filters; Holte, Denmark). The microphone was calibrated using a B&K Type 4237 sound pressure calibrator (94 dB at 1 kHz) to ensure accurate sound pressure measurements.

The scanning vibrometer, loudspeaker, and microphone were connected to a PSV-500 internal data acquisition board (Polytec PSV-500 vibrometer front-head; Waldbronn, Germany), which interfaced with a computer running Polytec 10.1.1 software for system control and data analysis. A broadband stimulus consisting of periodic chirps with a simulated frequency range of 2–10 kHz was delivered by the loudspeaker. To ensure consistent stimulus intensity, the amplitude of the broadband signal was mathematically corrected within the software, providing a uniform sound pressure level of 60 dB across all frequencies.

During measurements, the tympanic membrane region was selected within the software, and the laser focus was systematically adjusted to ensure high-quality data acquisition. The vibrations of the tympanic membrane were recorded in the frequency domain at a sampling frequency of 512 kHz. Data were post-processed and analysed using Polytec 10.1.1 software, and the final outputs were exported and visualised using Python libraries for graph plotting and detailed analysis.

### 2.6. Acoustic experiment on scaled acoustic trachea

The 20× scaled acoustic trachea was fixed in place using a clamp and stand system, with the model secured gently via a plastic clip to minimise mechanical interference. Acoustic measurements were conducted in an acoustically treated room using a combination of free-field and near-field stimuli.

A 1/8″ precision pressure microphone (Bruel & Kjaer, 4138; Nærum, Denmark), connected to a conditioning amplifier, was positioned adjacent to the spiracle opening to monitor and calibrate reference sound pressure levels during each trial. Two 25-mm probe microphones (Type 4182, Brüel & Kjær, Nærum, Denmark), each with a 25-mm-long and 1.24-mm inner diameter probe tube, were connected to preamplifiers (Type 2633, Brüel & Kjær). One probe was inserted into the spiracle opening, and the other at the distal end of the tracheal lumen, allowing simultaneous measurement of input and output sound pressures. All microphones were calibrated before testing using a B&K Type 4237 sound pressure calibrator (94 dB at 1 kHz) to ensure accurate and reproducible measurements. To acoustically isolate both ends of the trachea and prevent cross-contamination from airborne stimuli, a vertical barrier of acoustic foam was positioned between the spiracle and the open end of the model.

Two acoustic stimulation conditions were tested. In the near-field configuration (Fig. 4a), a custom probe speaker was positioned approximately 1 cm from the spiracle. This speaker was assembled by encasing an earbud in acoustic foam and attaching a plastic conical nozzle to direct the output into the spiracle opening, ensuring localised delivery of sound pressure. In the far-field configuration (Fig. 4b), a broadband loudspeaker (wide speaker) was placed 30 cm away from the spiracle. This setup simulates distant sound sources, where pressure waves arrive as planar fronts with relatively uniform phase and amplitude, analogous to natural bat calls in the wild.

**Fig. 4.**
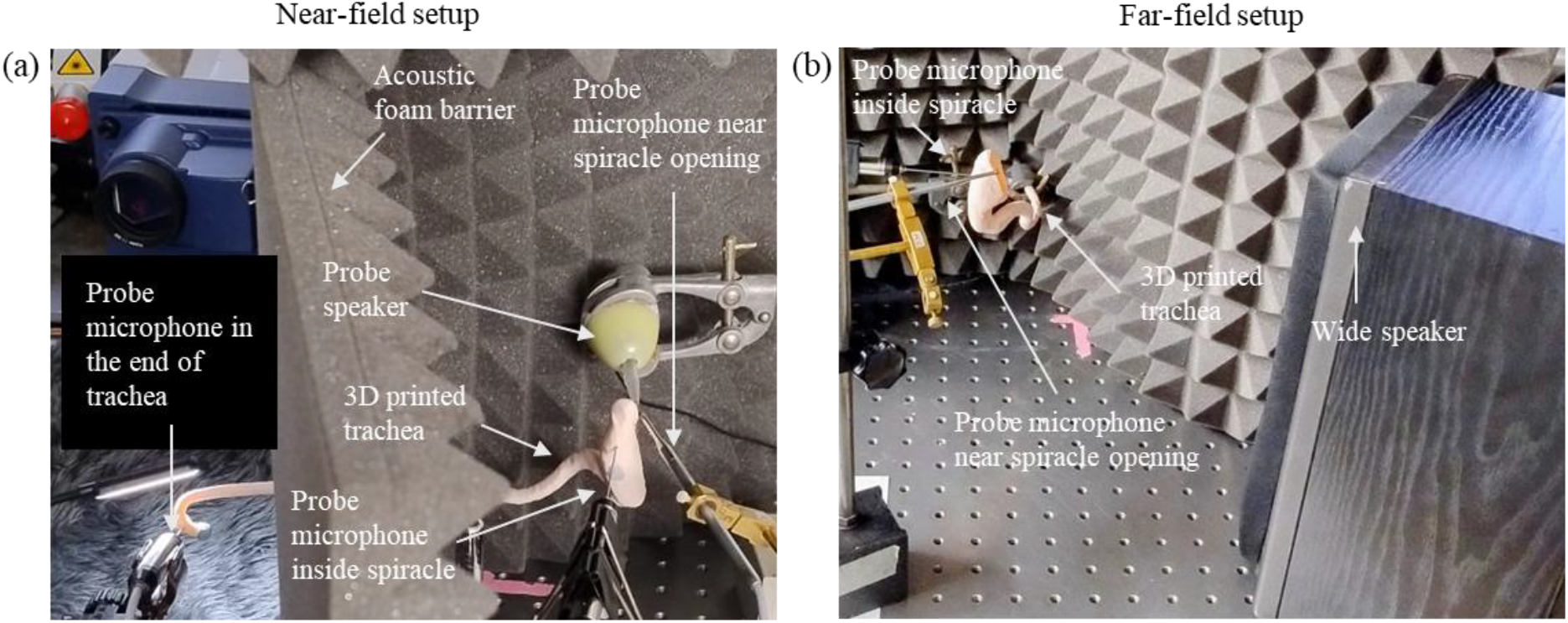
Acoustic characterisation setups for 3D-printed scaled katydid trachea. (a) Near-field configuration using a custom probe speaker positioned 1 cm from the spiracle to deliver focused acoustic input. Sound pressure was recorded using three probe microphones placed near the spiracle opening, inside the spiracle, and at the distal end of the trachea. An acoustic foam barrier was positioned to minimise cross-interference between the input and output ends. (b) Far-field configuration using a wide speaker placed 30 cm from the spiracle to simulate distant sound sources. Probe microphones recorded input and internal pressures, enabling comparison of acoustic transmission under different spatial stimulation regimes.

For both conditions, a broadband stimulus ranging from 0.5–4 kHz was delivered. To ensure consistent sound pressure across all frequencies, the stimulus amplitude was equalised within the presentation software using spectral correction, resulting in a flat 40 dB SPL broadband output.

## 3. Results

### 3.1. Displacement profile of real insect vs 3D printed TPU tympanic membrane

The displacement fields of a live *Copiphora gorgonensis* tympanum, with its pinna surgically removed and stimulated at 10 kHz from previous LDV study [9] and of the isolated 15 fold scaled TPU membrane driven at 7 kHz is displayed in Fig. 5. Spatial line profiles extracted across the tympana for both cases show negligible motion at the edges. Moving towards the centre, displacement rises steeply to a broad crest and then decays slightly asymmetrically for the real insect tympana toward the distal edge (Fig. 5a) and symmetrically for the 3D printed tympana(Fig. 5b). The living membrane reaches a peak of around 3.6 µm at roughly 200 µm from the proximal attachment, whereas the printed membrane peaks at around 0.9 mm at roughly 1.5 mm from the attachment.

**Fig. 5.**
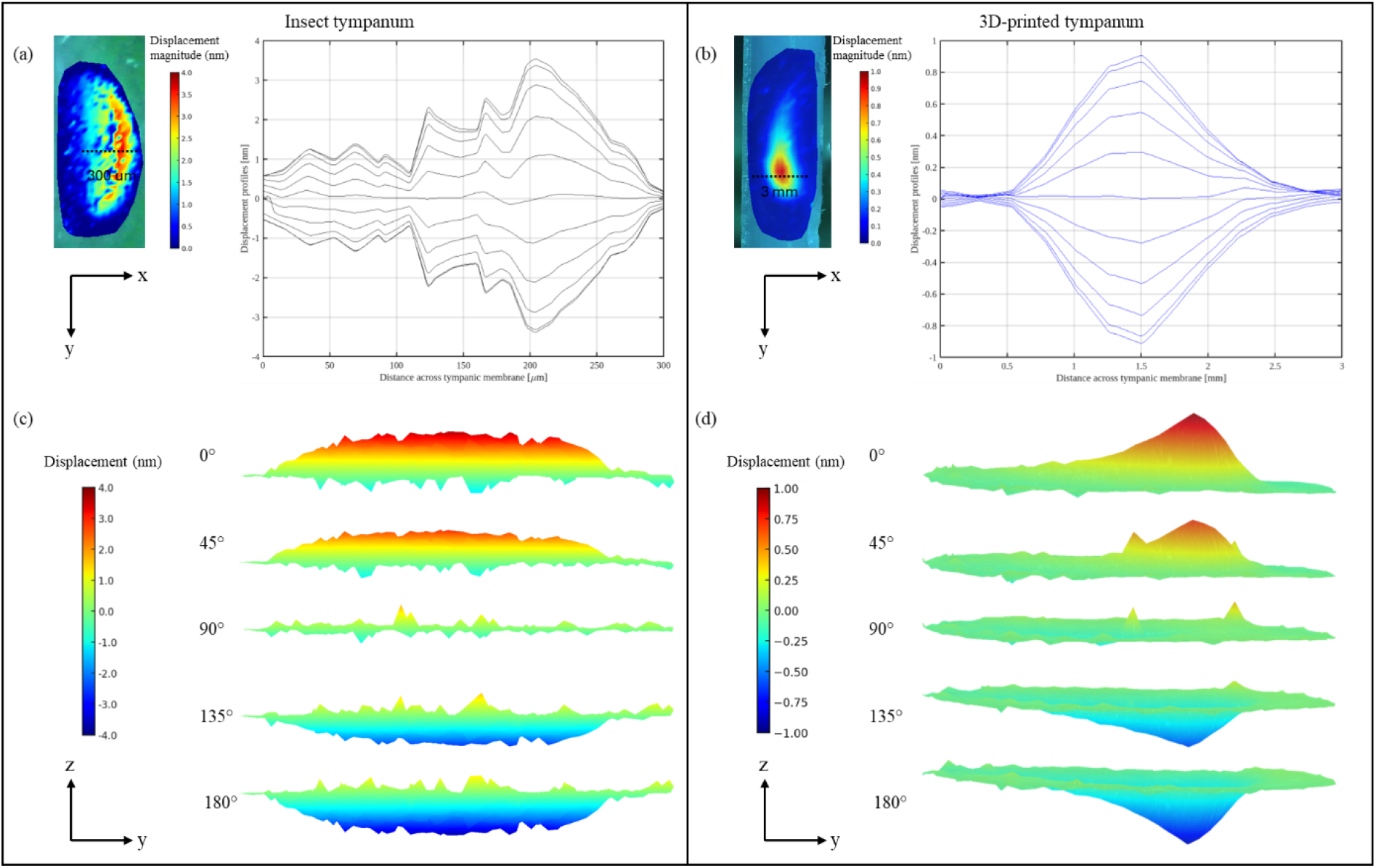
Displacement profiles of real insect and 3D printed isolated tympanic membrane. (a) LDV map of a live *Copiphora gorgonensis* tympanum at 10 kHz after pinna removal and the corresponding displacement profiles at different phases sampled along the dashed 300 µm transect. (b) LDV map of the 15 ×-scaled TPU membrane at 7 kHz with displacement profiles at different phases taken along the dashed 3 mm transect. (c) Sequential surface plots (five equally spaced phase snapshots within one stimulus cycle) of an *in-vivo Copiphora gorgonensis* tympanum with the pinna removed (10kHz excitation). The asymmetric displacement crest rises on the distal half of the membrane and collapses back through nodal regions toward the clamped rim. (d) Corresponding phase snapshots of the 15 ×-scaled TPU membrane driven at 7 kHz.

Five snapshots taken at equal phase intervals within a single acoustic cycle reveal that the live *C. gorgonensis* tympanum (pinna removed, 10 kHz stimulus) develops a broad, dome-shaped crest spanning across the membrane, with peak excursions of about ±4 nm, as presented in Fig. 5c. The complementary sequence for the 15 times larger TPU print (7 kHz; Fig. 5d) reveals a much narrower zone of activity: a sharp central ridge rises and falls while most of the surrounding film remains close to baseline, and the displacement range is limited to approximately +1 nm to –1 nm.

### 3.2. Laser-Doppler vibrometry of 3D printed assembled outer-ear structures

Complete pinna-tympanic membrane assemblies were fabricated in four material pairings: both PLA pinnae and membrane, both TPU pinnae and membrane, TPU pinnae with PLA membrane and PLA pinnae with TPU membrane (Fig. 6a). As the 3D-printed outer-ear models 15 times larger than the actual insect, the acoustic excitation was conducted in the 2–10 kHz range, corresponding to 30–150 kHz in the natural scale of bat echolocation calls. The recorded data were subsequently post-processed and frequency-scaled (Fig. 6b) for direct comparison of *C. gorgonensis* in vivo measurements [11]. The results revealed that assembly that employed the flexible TPU for the tympanic membrane exhibited substantial motion, displaying a broad resonance between 70–110 kHz. The PLA-pinna/TPU-membrane model produced a single displacement lobe with a peak amplitude of 1.93 nm at 75 kHz, whereas the all-TPU construct reached 1.27 nm at the same frequency. Both PLA-membrane combinations remained below 0.10 nm and showed no coherent spatial pattern.

**Fig. 6.**
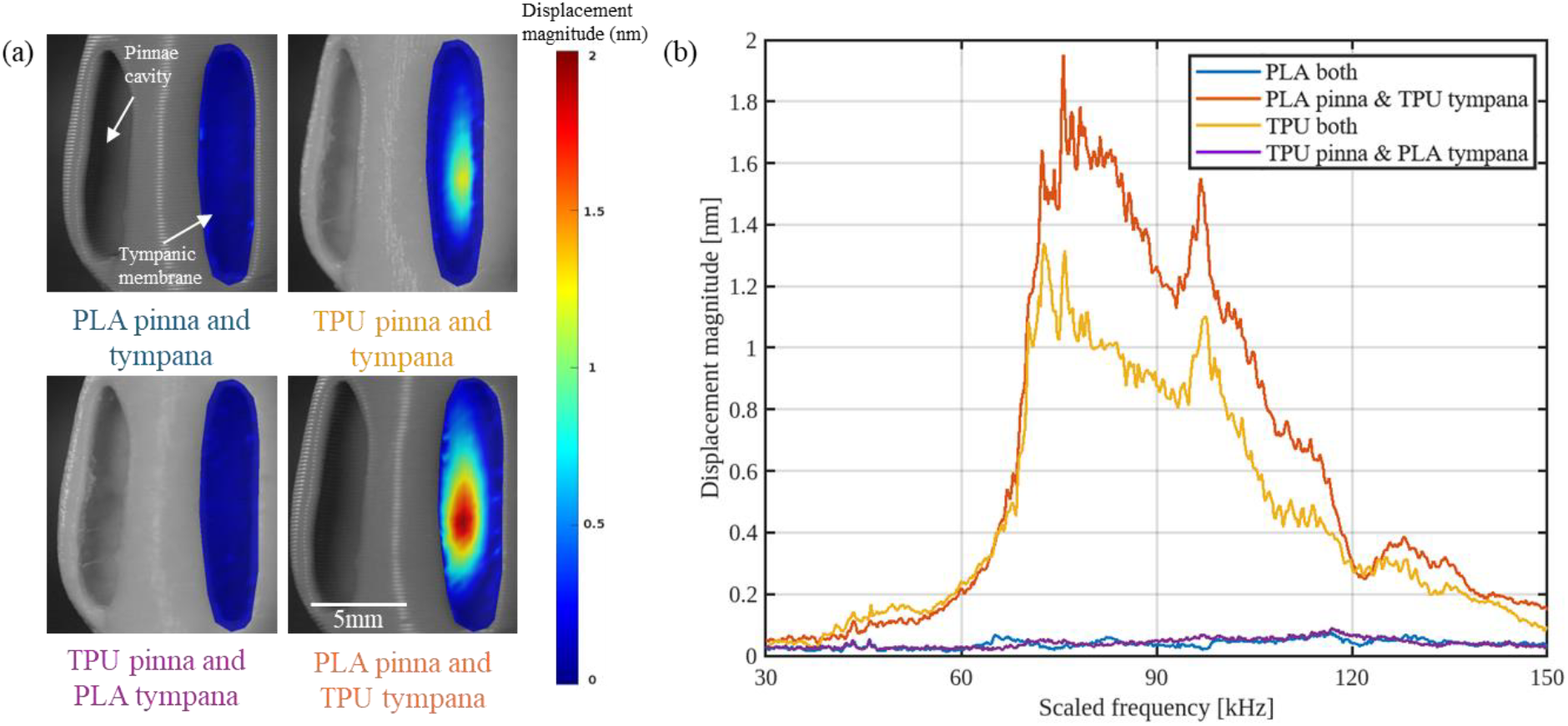
Pinna-membrane material pairing modulates tympanic vibration in assembled outer ears. (a) Displacement maps at a scaled 80 kHz for four pinna/membrane combinations. (b) Displacement magnitude at the membrane centre versus scaled frequency (30–150 kHz). The PLA-pinna/TPU-membrane assembly peaks at 1.93 nm, more than double the exposed-membrane value.

For the PLA-pinna/TPU-membrane construct, the transfer-function coherence between the acoustic stimulus and membrane displacement exceeded 0.95 from 60 kHz to ∼140 kHz and remained above 0.90 across the measured 30–150 kHz range (Fig. 7, red dashed line). The associated phase response (blue line) declined smoothly by ∼400° after 70 kHz, without abrupt jumps, indicating a largely linear, single-mode membrane motion. The high coherence confirms that most of the incident acoustic energy reached and drove the membrane throughout the frequency band examined.

**Fig. 7.**
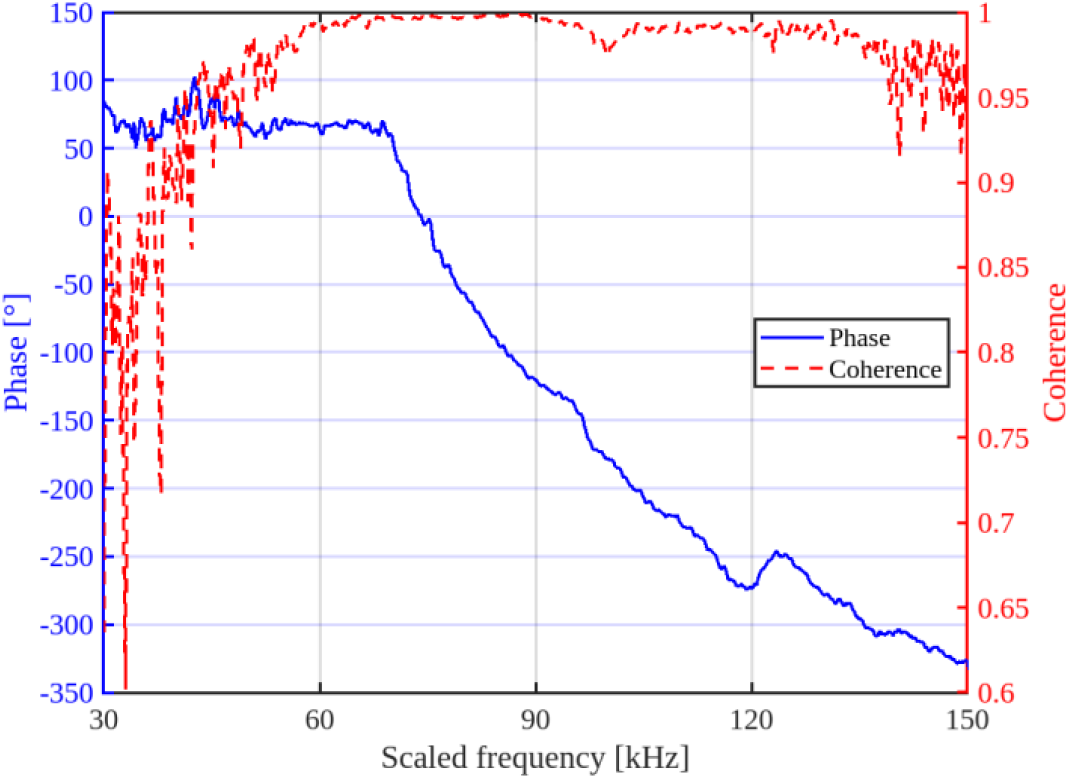
High-coherence, single-mode response of the PLA-pinna with TPU-membrane ear. Phase (blue) and transfer-function coherence (red) between the acoustic stimulus and membrane displacement across 30–150 kHz. Coherence exceeds 0.95 over most of the band, while phase decreases smoothly by ∼360°, indicating single-mode propagation.

### 3.3. Sound transmission through the 3-D-printed acoustic trachea

The 20 times scaled trachea, printed in PLA, was excited with broadband stimuli delivered either by a miniature probe speaker positioned 3 mm from the spiracle (“probe” condition) or by a 100 mm loudspeaker (wide speaker) placed 30 cm away (far-field condition) at 40 dB sound pressure level. As the tracheal model was enlarged 20 times, the acoustic experiments were conducted between 0.5–4 kHz, equivalent to 10–80 kHz in the actual insect. Pressure gain and unwrapped phase change were recorded at the spiracle opening and at the distal (tibial) end of the tube, and the results are shown in Fig. 8.

**Fig. 8.**
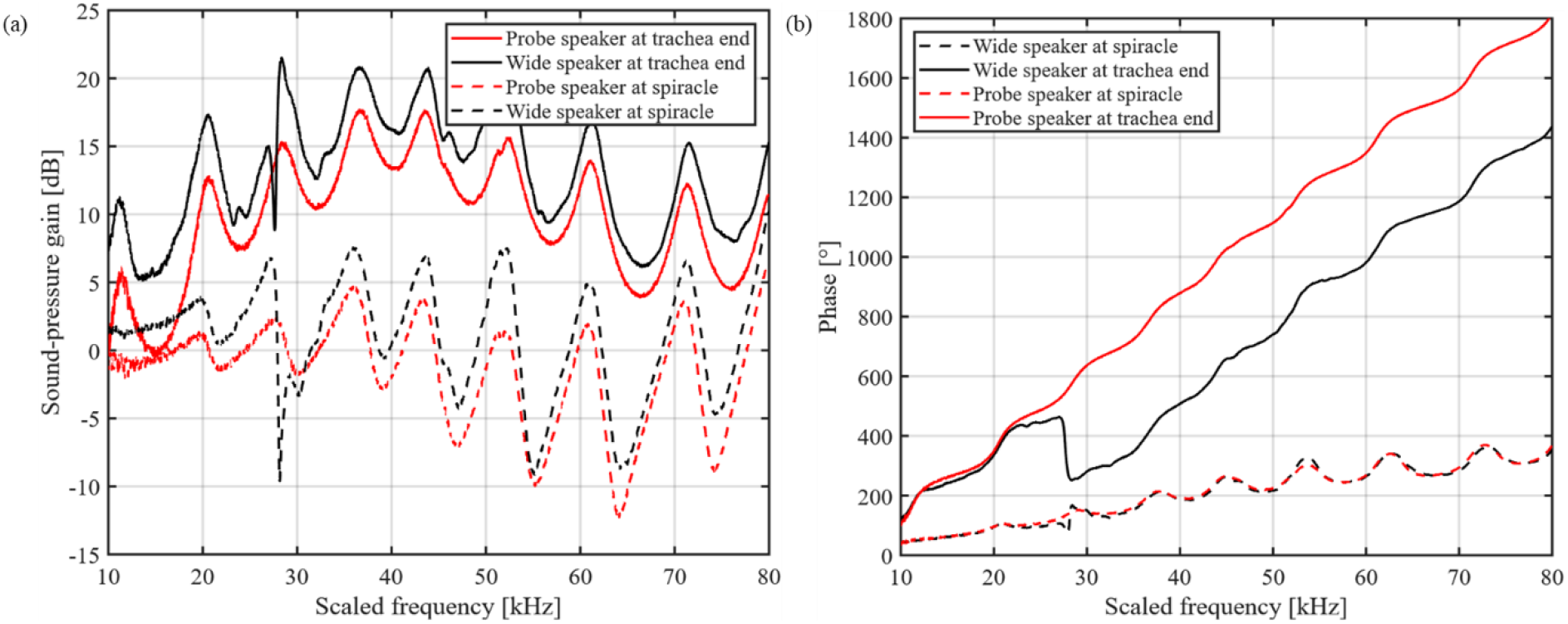
Sound-pressure gain and phase along the 3D-printed acoustic trachea. (a) Sound-pressure gain measured at the spiracle (dashed traces) and at the distal terminus (solid traces) for a near-field probe source positioned 3 mm from the spiracle (red) and for a far-field loud-speaker 30 cm away (black). Peaks of 17–21 dB at the terminus correspond to troughs at the spiracle, indicating that the opening acts as a frequency-selective filter while the exponential canal amplifies the transmitted bands. (b) Unwrapped phase for the same four conditions. Phase accumulates smoothly to ∼1,800° over the scaled 10–80 kHz sweep at the trachea end, confirming single-mode propagation along the horn; stimulus geometry alters phase gradient by <10 %.

Under far-field excitation (Fig. 8a, black curves), the tracheal end exhibited a series of broad gain maxima at ∼22, 32, 42, 52, 62 and 72 kHz, reaching 17–21 dB. At the spiracle, gain remained near 0 dB below 20 kHz and oscillated between −5 and +7 dB above that range. The near-field excitation (probe stimulus, red curves) produced the same spectral pattern but at slightly lower amplitude: peaks of 12–16 dB at the terminus and −6 to +5 dB at the spiracle. Thus, in both speaker configurations, the trachea increased sound pressure by more than an order of magnitude at its distal end across multiple ultrasonic bands.

The corresponding phase response (Fig. 8b) rose approximately linearly with frequency at the tracheal terminus, accumulating ∼1,700 ° between 10–80 kHz for the probe stimulus (red solid line) and ∼1,400 ° for the far-field stimulus (black solid line). The phase measured at the spiracle (dashed lines) remained below 350° over the same band and no abrupt discontinuities were observed, indicating a smooth propagation of the dominant acoustic mode along the tube. These measurements show that the printed horn preserves a frequency-dependent pressure build-up along its length while maintaining high coherence between input and output across the 10–80 kHz sweep.

## 4. Discussion

LDV revealed that the TPU film produced a single-lobe vibration pattern comparable to that of the living *Copiphora gorgonensis* exposed tympanum (see Supplementary Videos 1 and 2). Both displayed similar motion over a full acoustic cycle (Fig. 5), with the membrane rising on one side and settling back toward the edge. However, the live tympanum spreads motion over a broader area, whereas the print confines deflection to a sharper, localized peak. Also, the displacement height of the insect tympana reaches ±4 nm, while the scaled replica is limited to roughly ±1 nm. These discrepancies mainly come from the mismatch in material stiffness between the real insect tympanum and the printed TPU membrane. The insect’s tympanic membrane likely has a changing stiffness across its surface, which helps spread motion more evenly. In contrast, the TPU replica has a uniform modulus, so the vibration becomes concentrated in a smaller, sharper peak because of how the structure responds to sound. This difference in stiffness also explains why the live tympanum vibrates most strongly around 7 kHz, while the enlarged TPU model shows its main motion near 10 kHz, each resonating at its own natural frequency set by its material properties. Nonetheless, as the goal of this study is to detect tympanal vibrations through pinna cavity-induced resonance (which is influenced by structural shape than modulus) rather than matching the exact natural frequency of the membrane, these differences do not pose any significant issue.

LDV on assembled structures (Fig. 6) showed structures comprising of TPU membranes resonates in-between 70–110 kHz that closely aligns with the insect’s native bat-detection band (see supplementary video 3). This confirms that katydid pinnae behave as mechanical amplifiers for selective ultrasonic bands, a function previously inferred from LDV on intact legs [11]. The PLA pinna cavity with TPU membrane produced a 1.93 nm peak, 50 % larger than just TPU only. This confirms the acoustic properties of these structures being majorly influenced by architecture than just material properties like stiffness. Still, the PLA/TPU pairing captures the biological impedance contrast more faithfully, as the pinnae fabricated with PLA are orders of magnitude stiffer than TPU, reflecting most sound waves on tympana [29]. The higher stiffness PLA-membrane thus showed no sign of resonance. These findings are consistent with previous LDV data from real-ear measurements, which showed increased tympanic membrane vibration above 80 kHz in response to a broadband stimulus (20–120 kHz), with a resonant peak around 107–111 kHz attributed to pinnae-driven amplification [11]. Phase fell gradually across the measured band, indicating essentially direct pressure loading with no abrupt modal transitions. The assembled PLA-pinna/TPU-membrane model exhibited coherence >0.95 throughout the frequency band (Fig. 7), confirming that nearly all incident acoustic energy was transmitted to the membrane.

Pressure mapping along the 20 times scaled PLA acoustic trachea revealed that gain at the spiracle oscillates between −5 dB and +7 dB, whereas gain at the distal tip reaches 17–21 dB (Fig. 8a). The oscillating sound-pressure gain indicate that the spiracle operates as a frequency-selective filter, while the exponential canal functions as a passive amplifier [30]. Phase advanced smoothly to ∼1,700 ° over 10–80 kHz (Fig. 8b), matching a similar profile measured in intact insect trachea. The phase shift at 23 kHz was approximately 460° (around 1.3 cycles), same as the real insect tracheal phase shift (460–490° or 1.3–1.4 cycles) [9], supporting the consistency of 3D printed model for experimentation and analysis. Changing from a distant far-field loudspeaker to a near-field probe altered peak gain by <5 dB and phase by <10 %, suggesting that the horn’s filtering properties are largely source-independent.

Superimposing the measured these sound-pressure gains onto the finite-element study [17] shows peaks aligning within ±2 kHz and amplitudes within ±3 dB across 10–80 kHz (Fig. 9). This agreement validates the 3D printed scale model, providing the first physical confirmation of that model’s predictions. Thus, printed horn can be instrumented at arbitrary points and offers a powerful benchmark for refining future simulations.

**Fig. 9.**
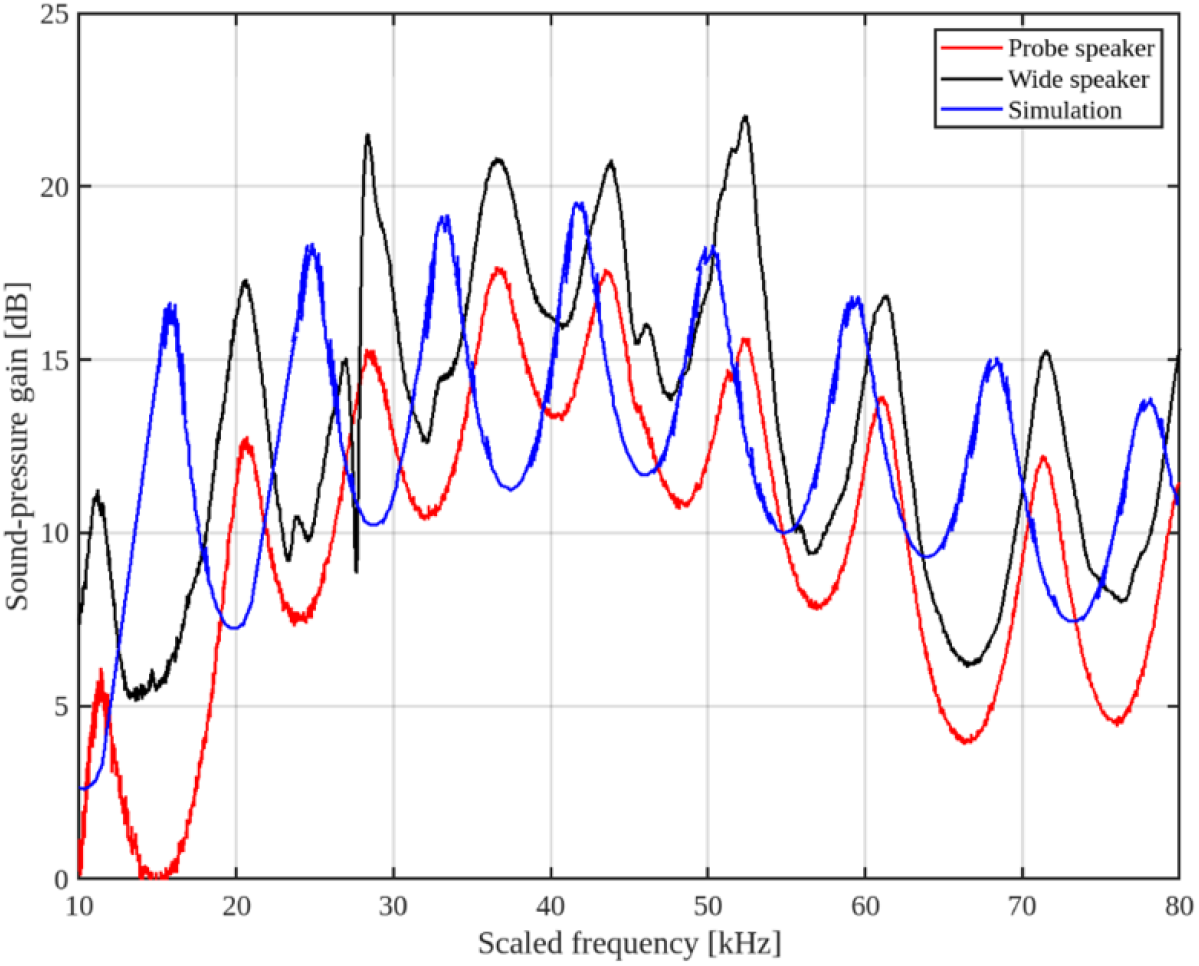
Printed tracheal gain closely matches finite-element simulation. Sound-pressure gain at the tracheal terminus measured with probe (red) and wide (black) sources compared with the finite-element prediction (blue) over a scaled 10–80 kHz range; peaks align within ±2 kHz and ±3 dB.

## 5. Conclusion

The scaled 3D-printed models of the *Copiphora gorgonensis* outer ear successfully reproduced the biological hearing function observed in the living insect. The combination of flexible membranes and rigid pinnae, operated as a natural bat detector, showing mechanical gain within the ultrasonic echolocation band of bat calls. Also, the printed acoustic trachea exhibited the filtering and amplification behaviour, previously predicted only by numerical simulations, confirming that the exponential horn acts as a passive amplifier and the spiracle as a filter for frequency-selective input.

The process of producing scaled physical models provides a reliable experimental platform for testing hypotheses about miniature pressure-difference receivers while reducing the need for live specimens. The controlled and repeatable fabrication with design freedom of 3D printing process enables allows future exploration of other key parameters such as influence of stiffness, cavity geometry, and horn shape. This provides a powerful tool for acoustic biophysics and while also aligning with the 3Rs principles by working on models. Importantly, this interdisciplinary approach demonstrates that insect-inspired hearing mechanisms can be translated into scalable miniature engineering systems.

The validated relationships between cavity and trachea structure, resonance, and phase delay act as a foundation for developing miniaturised mechanical adapters, MEMS-based microphones, and bio-inspired directional sensors that exploit passive acoustic filtering and amplification and reducing dependence on electronic gain and software post-processing, thus increasing energy efficiency. As 3D printing technological advance, capable of printing higher resolution and integrate multi-material naturally, replicating these biological structures in actual scale will become feasible. In the future, this approach will allow us to combine the outer-ear and acoustic trachea models into a single integrated structure. Such a unified model will enable direct experimentation on the katydid’s dual-input hearing mechanism, providing deeper insight into its acoustic properties and further bridging biological understanding with technological innovation through tangible, nature-inspired design.

## Supporting information

Real insect tympanic membrane motion

Replicated TPU Tympana motion

Assembled PLA cuticle & TPU Tympana motion

## Acknowledgements

This research is part of the project “Biophysical and ecological function of microscale ears using scaled 3D prints” funded by the Leverhulme Trust [Grant RPG-2023-204] to Fernando Montealegre-Z; and was also funded by the Natural Environment Research Council (NERC), [grant DEB-1937815] to Fernando Montealegre-Z.

## CRediT authorship contribution statement

**Md Niamul Islam:** Conceptualization, Methodology, Software, Validation, Formal analysis, Investigation, Data curation, Visualisation, Writing – original draft, Writing – review & editing.

**Fabio A. Sarria-S:** Methodology, Software, Validation, Investigation, Data curation, Visualisation, Writing – review & editing.

**Fernando Montealegre-Z:** Conceptualization, Resources, Writing – review & editing, Supervision, Funding acquisition, Project administration.

